# FRaeppli, a multispectral imaging toolbox for cell tracing and dense tissue analysis in zebrafish

**DOI:** 10.1101/2022.01.14.476353

**Authors:** Sara Caviglia, Iris A. Unterweger, Akvilė Gasiūnaitė, Alexandre E. Vanoosthuyse, Francesco Cutrale, Le A. Trinh, Scott E. Fraser, Stephan C. F. Neuhauss, Elke A. Ober

**Author notes:** equal contribution (Sara Caviglia, Iris A. Unterweger).

## Abstract

Visualizing cell shapes, interactions and lineages of differentiating cells is instrumental for understanding organ development and repair. Across species, strategies for stochastic multicolour labelling have greatly facilitated tracking cells in *in vivo* and mapping neuronal connectivity. Nevertheless, integrating multi-fluorophore information into the context of developing tissues in zebrafish is challenging given their cytoplasmic localization and spectral incompatibility with commonly used fluorescent markers. Here, we developed FRaeppli (<u>F</u>ish-<u>Raeppli</u>) expressing bright membrane-or nuclear-targeted fluorescent proteins for efficient cell shape analysis and tracking. High spatiotemporal activation flexibility is provided by the Gal4/UAS system together with Cre/lox and/or PhiC31integrase. The distinct spectra of the FRaeppli fluorescent proteins allow simultaneous imaging with GFP and infrared subcellular reporters or tissue landmarks. By tailoring hyperspectral protocols for time-efficient acquisition, we demonstrate FRaeppli’s suitability for *live* imaging of complex internal organs, like the liver. Combining FRaeppli with polarity markers revealed previously unknown canalicular topologies between differentiating hepatocytes, reminiscent of the mammalian liver, suggesting shared developmental mechanisms. The multispectral FRaeppli toolbox thus enables the comprehensive analysis of intricate cellular morphologies, topologies and tissue lineages at single-cell resolution in zebrafish.

## Introduction

A prerequisite for understanding how distinct tissues form is the capability to visualize the resident cell types at single-cell resolution (Megason and Fraser, 2007). This information is essential for uncovering the cellular architecture of organs and entire embryos, as well as the intricate cellular processes, such as cell interaction, movement and proliferation, guiding tissue formation. The transparency and rapid growth of zebrafish are ideal for dissecting the cellular processes establishing tissue architectures in vertebrate development and disease. Uniform membrane labelling at the tissue level generates an approximate view of cellular shapes and is therefore suitable for uniformly-shaped cells, while elaborate cellular morphologies and intricate contacts, for instance via protrusions (Caviglia and Ober, 2018), remain elusive. Instead, sparse labelling by expression of a single membrane-targeted fluorescent protein (FP) (e.g. (Cayuso et al., 2016; Sanders et al., 2013) outlines individual cells and has significantly advanced our understanding of dynamic cell behaviours. Nevertheless, information concerning the cellular topologies, the number and morphology of contacts between adjacent cells and the nature of their interactions is frequently lacking, because the cellular microenvironment remains unlabelled.

The need to distinguish neighbouring cells has been met partially by the development of the transgenic *Brainbow* system (Livet et al., 2007), which randomly labels neighbouring cells within a population by the stochastic expression of three or four FPs (reviewed in (Richier and Salecker, 2015; Weissman and Pan, 2015). Stochastic combinatorial expression of spectrally distinct FPs is achieved by their sequential arrangement in a DNA cassette, with each FP flanked by recombinase recognition sites, permitting stochastic recombinase-mediated DNA excision or inversion. This approach, pioneered in mice, subsequently has been adapted in a variety of species, including zebrafish (Livet et al., 2007) (Gupta and Poss, 2012; Pan et al., 2013; Richier and Salecker, 2015; Weissman and Pan, 2015), enabling key advances ranging from uncovering distinct neural networks (e.g. (Fontenas and Kucenas, 2021; Forster et al., 2017) to clonal dominance of stem cell-like populations in differentiating tissues (e.g. (Gupta and Poss, 2012; Gurevich et al., 2016). Brainbow-tools targeting FPs to specific cellular compartments, such as the membrane, have been successfully used in mice and flies, but only rarely in zebrafish (Forster et al., 2017). Moreover, comprehensively resolving the complex organization of a developing tissue requires additional information of specific cell types, subcellular structures (e.g. cytoskeleton) or signalling pathway activity as landmarks. A multitude of experimental possibilities will therefore open up when multicolour cell labelling systems are combined with spectrally distinct signals, such as infrared proteins (Cook et al., 2019). However, the majority of existing reporter lines marking tissues and cell types express GFP, which overlaps with the spectra of existing Brainbow-like systems and thereby preclude their combined use.

To permit more distinct labelling of cells, we introduce here the FRaeppli (<u>F</u>ish <u>Raeppli</u>) tools, transgenic lines inspired by Raeppli, a whole-tissue labelling system developed in *Drosophila melanogaster* (Kanca et al., 2014). FRaeppli expresses four instead of three bright monomeric FPs: mOrange, mKate2, mTFP1 and TagBFP, which are spectrally distinct from GFP and far red and infrared fluorophores, allowing for the simultaneous acquisition of six distinct labels. We demonstrate the versatility of the membrane-tethered *fraeppli-caax* lines for large-scale visualization of cell topologies and the nuclear *fraeppli-nls* for tracing of cell lineages in dense tissues. We show its versatility with examples of larval and adult tissues, genetic strategies to control clone size, *live* imaging and a hyperspectral protocol for time-efficient data acquisition.

## Results/Discussion

### Design and general expression characteristics of FRaeppli

For a versatile next generation multicolour cell labelling system in zebrafish, we adapted *Drosophila* Raeppli (Kanca et al., 2014) and produced <u>F</u>(ish)<u>Raeppli</u>. The colour cassette consists of four bright fluorescent proteins (FPs): mOrange (E2-Orange in *fraeppli-nls*), mKate2, mTFP1 and TagBFP (Figs. 1A, S1). Based on their excitation and emission spectra, distinct imaging of these FPs can be performed with fluorescent molecules in the green/GFP and far-red range. FRaeppli is therefore compatible with a wide variety of established transgenic lines as well as diverse cell-and organelle labels. We generated two different versions for application-specific use: *fraeppli-nls* targets the FPs to the nucleus with an added nuclear localization signal (NLS; Fig. 1B,C) for cell lineage and growth analyses, whereas *fraeppli-caax* directs the FPs to the cell membrane with a Ras farnesylation sequence (CAAX) for cell shape analyses (Fig. 1E). Recombination of the colour cassette is mediated by the bacteriophage PhiC31 integrase (Bischof et al., 2007; Lister, 2011; Thorpe et al., 2000), so far only utilised for transgenesis in zebrafish (Hu et al., 2011; Mosimann et al., 2013). PhiC31 integrase catalyses efficient recombination of unidirectional *attB* and *attP* sites, excising the DNA fragment flanked by these sites. In FRaeppli, a single *attB* site is placed directly after the promoter and *attP* sites in front of each of the four FP open reading frames (Fig. 1A). Recombination events result in new *attL* and *attR* sites, incompatible with further PhiC31-mediated recombination and thereby initiating stable expression of one of the four FPs per cell (Fig. S1). PhiC31 integrase is built into the FRaeppli cassette and controlled by an upstream 1,5 kb STOP-cassette flanked by two *lox2272* sites. Cre-mediated recombination of the two *lox2272* sites excises the STOP-cassette, placing PhiC31 integrase expression under the direct control of the regulatory *Upstream Activating Sequence (UAS)*. The expression of the FRaeppli construct is driven by the binary Gal4/UAS system (Brand and Perrimon, 1993). Binding of the transcriptional activator Gal4 or optimized versions, such as KalTA4 (Distel et al., 2009), to the regulatory *UAS* elements drives the transcription of both the PhiC31 integrase and the randomly selected FP, following integrase-mediated *attB/attP* recombination (Fig. S1D).

**Fig. 1.**
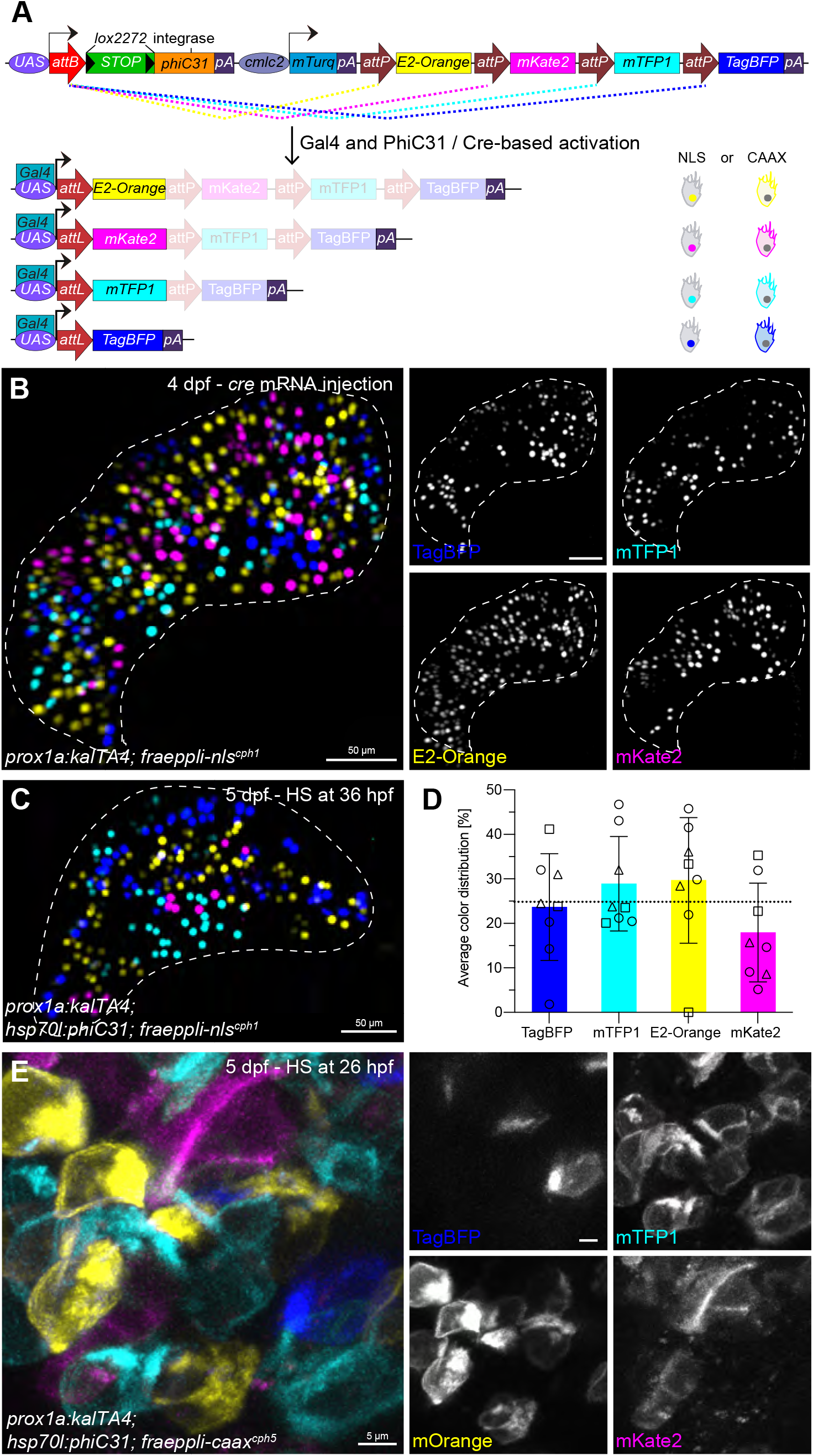
FRaeppli expression is controlled by sequential Gal4 and Cre/PhiC31 integrase activity. (A) Schematic of FRaeppli activation. Cre catalyses excision of a stop cassette, priming FRaeppli for activation. Gal4/UAS initiates *phiC31 integrase* expression, which drives *attB* and *attP* recombination and stochastic expression of one out of four FPs. FPs are targeted to the cell membrane in *fraeppli-caax*, or to the nucleus in *fraeppli-nls*. The FP order represents *fraeppli-nls*; in *fraeppli-caax* it is mKate2-mTFP1-mOrange-TagBFP; Fig. S3E). (B, C) Activation of *fraeppli-nls; prox1:kalTA4* by *cre* mRNA injection (B) or *hsp70l:phiC31* (C) results in clones with nuclei in all four colours. (D) Quantification of the colour selection frequency of *fraeppli-nls*; N=3, n=8, Error bars represent SDs. (E) Activation of *fraeppli-caax; prox1a:kalTA4* with *hsp70l: phiC31* generates clones outlining cells with all FRaeppli colours.

We tested FRaeppli recombination in the forming liver first by injecting *cre* mRNA into stable transgenic *fraeppli*-*nls* or *fraeppli*-caax embryos, leading to clones in four colours in the majority of embryos (Figs. 1B, S2A). This indicates the correct expression of the FRaeppli *phiC31 integrase* and the subsequent induction of all four possible recombination events. Next, we tested recombination after conditional Cre expression by crossing *fabp10a:kalTA4; fraeppli-caax* to *hsp70l:cre* and subjecting embryos to a heat-shock at 33 hours post fertilisation (hpf). They expressed all four FPs in the liver (Fig. S2B,C), indicating that all functional elements of FRaeppli work correctly. Three independent lines each of *fraeppli-nls* and *fraeppli-caax* were identified, *fraeppli-nls*^*cph1-cph3*^ and *fraeppli-caax*^*cph4-cph6*^, with different levels of transgene activation (Fig. S2A-D). Most experiments were performed with the most penetrant lines: *fraeppli-nls*^*cph1-cph3*^ and *fraeppli-caax*^*cph4*^.

We expanded the activation strategies for conditional FRaeppli labelling by generating an inducible transgenic PhiC31 integrase line, *hsp70l:phiC31*^*cph7*^. This strategy is faster, as it bypasses Cre-induced excision of the STOP cassette and directly activates recombination and colour selection therefore more suitable also for early stages of embryogenesis (Fig. S1C). Similar to the FRaeppli-inbuilt PhiC31 integrase, activation of exogenous PhiC31 integrase expression in *prox1a:kalTA4; hsp70l:phi*^*cph7*^; *fraeppli-nls* or *fraeppli-caax* larvae efficiently induces all four recombination events (Fig.1 C,E). Examining mRNA and protein expression over time, by in situ hybridization chain reaction (HCR; (Choi et al., 2018) and *hsp70l:phiC31-sfGFP-NLS* expression, respectively, showed increasing expression of *phiC31* mRNA between 2-4 hours and of Phi31 protein 4 hours post heat-shock (Fig. S3A-C; movie S1). Direct induction of FRaeppli colour recombination by PhiC31 integrase allows subsequent temporally separated Cre-mediated gene inactivation, impossible for Cre-controlled multicolour tools.

Unbiased colour selection ensures the differential labelling of neighbouring cells. We therefore assessed colour representation using different transgene activation strategies and determined no significant preference for one or more colours in *fraeppli-nls* or *fraeppli-caax* (Figs. 1D, S2E). Nonetheless, we noticed that the timely appearance of the FPs can vary, mostly starting with tagBFP-CAAX or tagBFP-NLS (Fig. S3D), followed by the other FPs and mKate2-CAAX or E2-Orange-NLS appearing last. Depending on the experimental context, the lag time between the first and last FP can take between 10-30 hours. To exclude that this could be due to an asynchronous PhiC31 Integrase-mediated recombination, we assessed recombination of the colour cassette at the genomic level (Fig. S3E-G) performing PCR amplification at 10 and 24 hpf in single *hsp70l:gal4; fraeppli-caax*^*cph4*^ embryos injected with *phiC31 integrase* mRNA. This showed successful recombination of all four FP loci in individual embryos already at 10 hpf (Fig. S3F-F’’’), long before any colour was detectable, indicating a synchronized recombination during blastula stages. Likewise, *cre* mRNA-injected embryos of the same genotype show efficient amplification of all recombined FP loci already 24 hours after heat-shock, indicating that FRaeppli-inbuilt PhiC31 Integrase can also drive the stochastic recombination of all FP loci (Fig. S3G). The lag in FP appearance is likely due to a combination of individual FP maturation times, for instance 13 min for tagBFP compared to 78 min for E2-Orange (www.fpbase.org) and mRNA stability issues due to the presence of a single polyA in the colour cassette, after tagBFP. The farther the respective FP is from the polyA the longer it takes to build up detectable FP protein levels. We further noticed that the strength of a given Gal4 driver influences the temporal gap between recombination and colour detection. For instance, in the liver it can take between 14-30 hours for all FRaeppli FPs to appear, after stimulating recombination using *fabp10a:kalTA4* or *prox1a:kalTA4* respectively. Experimental design should therefore consider the strength of the Gal4 driver with respect to the time of colour detection. These results corroborate that the FRaeppli lines stochastically label cells by simultaneous recombination events, pivotal for tissue morphology and lineage tracing studies.

### Temporal and spatial activation of FRaeppli cell labelling

A key feature of multicolour cell labelling is the versatile control of recombination in space and time, for instance to mark progenitor cells before the onset of differentiation for lineage tracing or labelling of individual cells or small clones, essential for high-resolution cell morphology and cell interaction studies. This can be achieved by labelling cell populations of choice at defined time-points preceding analysis. Different PhiC31 Integrase, Cre and Gal4 expression methods were assessed for varying clone size by temporally controlling FRaeppli recombination in the liver (Fig 2G). Large clones and dense labelling can be achieved by *phiC31 integrase* mRNA injection mediating recombination during blastula stages and *prox1a:kalTA4* induced FP expression in liver progenitors (Fig. 2A). Clones are slightly smaller when *cre* mRNA injection removes the STOP cassette and *prox1a:kalTA4* controls recombination by subsequent expression of the FRaeppli-inbuilt PhiC31 Integrase (Fig. 2B). Late induction of recombination by *hsp70l:cre* at 70 hpf results in small clones and lower labelling density (Fig. 2C). Similar activation strategies lead to progressively smaller clones using the hepatocyte driver *fabp10a:kalTA4* (Fig. 2D-F), demonstrating that different tissue-specific drivers can control FRaeppli expression, as well as the accessibility of FRaeppli for recombination at later developmental stages. The combination of the Gal4/UAS and Cre/lox systems opens up the possibility to select a subset of cells within a population by the use of two different promoters. For instance, this is achieved by intersection of a tissue-wide Gal4 driver with a cell type-specific Cre- or CreERT2 line or vice versa. This dual activation strategy enables the analysis of new subpopulations arising from genome-wide transcriptome studies by visualizing cell shapes, behaviours and lineage at single cell resolution.

**Fig. 2.**
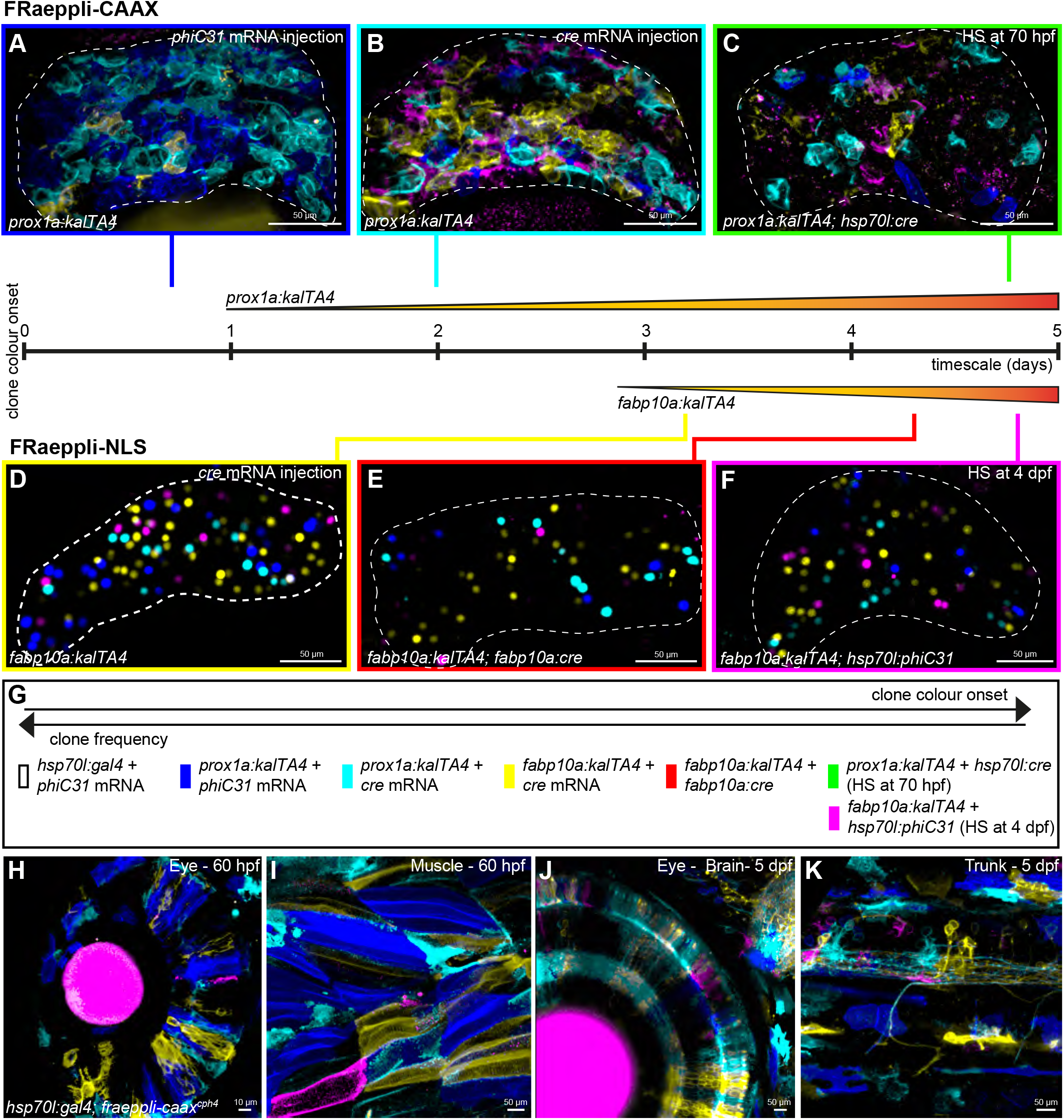
Versatility of spatiotemporal FRaeppli activation. Initiating FRaeppli recombination by *phiC31 integrase* mRNA injection (A) results in larger and denser clones than with *cre* mRNA injection (B, D). Late activation with inducible (C) or cell-type specific transgenic *cre* lines (E, F) results in smaller clones. Onset of the expression of the Gal4 driver also influences the temporal appearance of clones (see timeline in G). (H-K) Widespread FRAeppli-CAAX expression in tissues throughout the embryo, following injections with *phiC31* (H-I) or *cre* mRNA (J-K). Transgene expression was maximized by daily heat-shocks starting at 26 hpf, until imaging. 60 hpf (H,I) and 5 dpf (J,K) larvae are shown.

To validate FRaeppli for broad labelling in the embryo, we employed the ubiquitous driver *hsp70l:gal4* and *phiC31 integrase* or *cre* mRNA injection in *fraeppli-caax*^*cph4*^ embryos. Clones of varying size can be achieved by initiating recombination through induction of *gal4* expression at different time-points. Early induction at 26 hpf resulted in large clones spread throughout the body (Fig. 2H,I); later induction at 50 hpf also led to broad variegated labelling, though of smaller clone size (Fig. 2J,K). Thus, FRaeppli is a highly versatile cell labelling system, as multiple approaches can be employed to control temporal and broad or tissue-specific multicolour cell labelling, which is achieved by the specific combination the Gal4/UAS, Cre/lox and PhiC31 integrase systems built into FRaeppli.

### Advanced image acquisition and cell labelling strategies for FRaeppli

One challenge for imaging multispectral cell labelling is the frequently overlapping excitation and emission spectra of the used FPs and fluorophores (Cutrale et al., 2017; Dickinson et al., 2001). This can be addressed by sequential imaging to separate the four FRaeppli FPs (Fig. S4A). However, imaging time increases concomitantly with the number of colours (Fig. S4B). Alternatively, simultaneous excitation and acquisition of all channels by spectral imaging reduces imaging time (Dickinson et al., 2001). First, we acquired all spectral data of larval *prox1a:kalTA4*; *fraeppli-nls* livers by lambda scanning (Fig. 3A), using 32 channels of 8,9 nm width, thereby collecting signal from the 410-694 nm range, effectively reducing imaging time by 2-4 fold. Second, we employed the Hyper-Spectral Phasor software (HySP), which transforms the acquired spectral information by Fourier transformation into a 2D PhasorPlot, (Fig. 3A’) for subsequent label-unmixing by defining distinct spectral signatures (Fig. 3A’’; (Cutrale et al., 2017). This powerful method allows fast unmixing of large spectral images and without the need for calibration with reference samples for each spectral signature. Directly comparing the same *prox1:kalTA4*; *fraeppli-nls* liver acquired sequentially with band pass filters (Fig. S4D) or spectrally (Fig. S4E) showed that hyperspectral imaging combined with HySP processing provides additional spectral separation of noise derived from auto-fluorescence (Cutrale et al., 2017), advantageous for metabolically active tissues such as the liver. This method can be applied to *live* and fixed tissues of *fraeppli* larvae as well as adults (Figs. 3D, S4C,E).

**Fig. 3.**
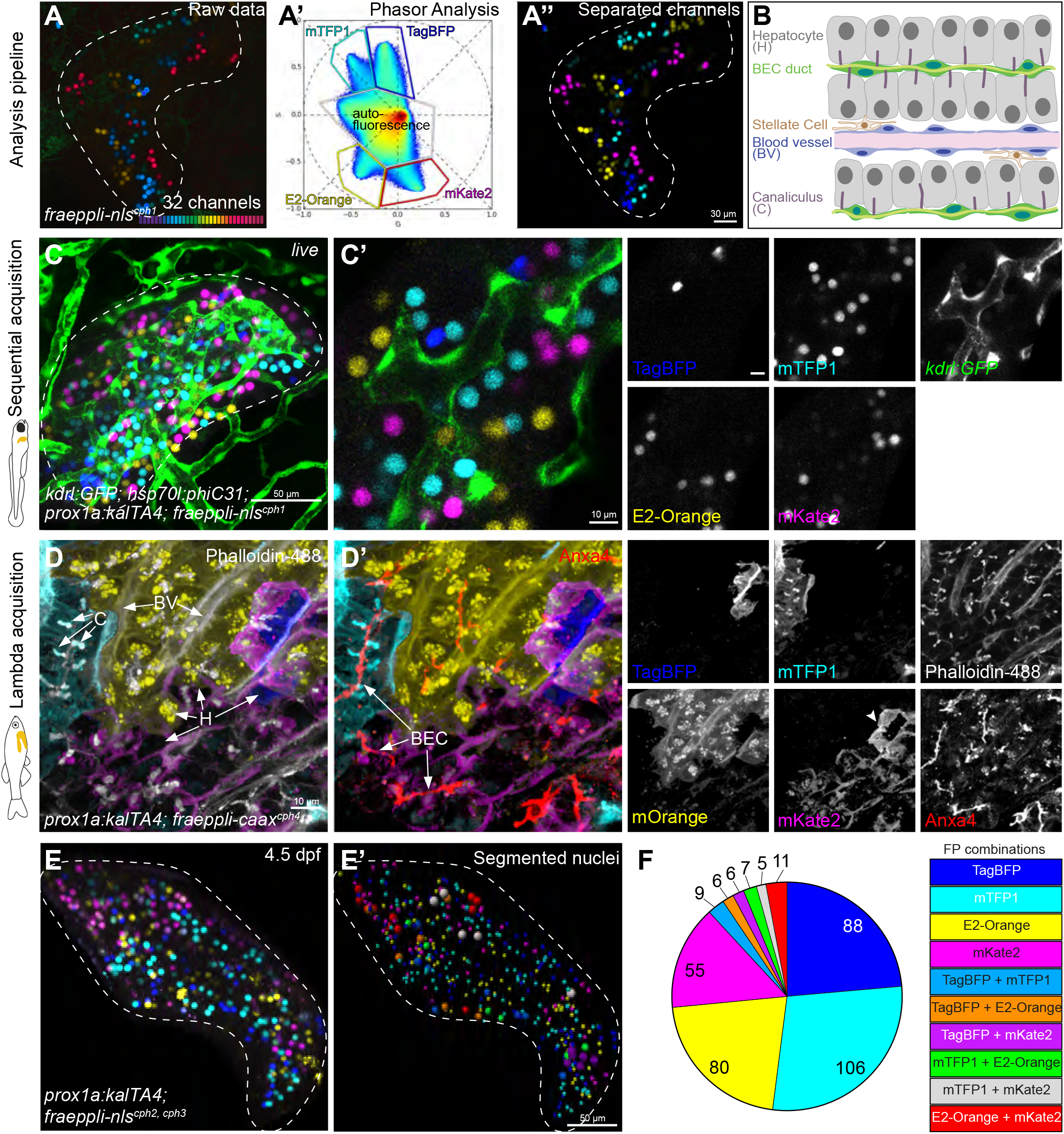
Advanced strategies for FRaeppli acquisition and multiplexed tissue labelling. (A) Spectral imaging of FRaeppli (A’) can be unmixed by hyperspectral phasor analysis (A”) into four separate channels (A’’’). (B) Schematic showing the functional zebrafish liver architecture. (C, C’) FRaeppli + 1 label: *Live* imaging of 5 dpf liver co-expressing hepatic FRaeppli-NLS and endothelial transgene *kdrl:GFP* highlighting vascular network. (D,D’) FRaeppli + 2 labels: Spectral acquisition of one year-old adult liver stained for Phalloidin (green) outlining liver architecture (D) and simultaneously for Anxa4 (red) to visualize biliary epithelial cell (BEC) ducts (D’). (E-F) Combination of two *fraeppli-nls* transgenes generates clones expressing ten discernible hues in the 5 dpf liver (E). Nuclei were segmented and pseudo-coloured for analysis, with nuclei expressing two different FPs have a larger diameter for visualization (E’). Absolute frequencies of different colour combinations in the liver (F).

Adding cellular and subcellular markers to FRaeppli-labelled populations would provide important tissue-specific landmarks and significantly extend the experimental possibilities for morphology and lineage studies. To achieve this, we tested the compatibility of FRaeppli with cell type-specific labels marking subpopulations in distinct spectra. First we combined *fraeppli-nls* with the endothelial *kdrl:gfp* line, showing *live* how hepatic clones arrange along the blood vessel network and the clear separation of all labels, including GFP (Fig. 3CC’). Next, we visualized the liver architecture (Fig. 3B) staining *fraeppli-caax* and *fraeppli-nls* adult liver sections and larval livers for Actin and Anxa4 or TO-PRO, highlighting the hepatocyte canaliculi, blood vessel network and biliary network or nuclei, respectively (Figs. 3D-D’, S5A-B’). These applications demonstrate the compatibility of FRaeppli with at least two spectrally distinct labels, enabling simultaneous clonal labelling with landmarks at the tissue, cellular and subcellular scale. Notable, FRaeppli expression is not only stably maintained into adulthood, but also suitable for SeeDB2 clearing (Ke et al., 2016) (Figs. S5A; movie S2), permitting long-term tracing and clonal analysis.

Moreover, elevating the number of distinct colour hues, for differential labelling of cells within a population can be achieved by inducing colour recombination in embryos carrying multiple transgene copies. Crossing two *fraeppli-nls* alleles demonstrates an increased colour diversity in the developing liver up to ten colours (Fig. 3E,E’), indicating successful recombination of all possible colour combinations, with comparable cell numbers expressing two distinct FPs (Fig. 3F). Altogether, we established the compatibility of FRaeppli with hyperspectral imaging and analysis strategies, tissue clearing approaches and common GFP and infrared tools significantly expanding experimental possibilities for high-resolution multicolour morphological and functional studies.

### Deciphering complex tissue organization: using FRaeppli to study hepatocyte polarity and interactions in the zebrafish liver

A powerful advantage of FRaeppli-CAAX is to differentially label the morphologies of adjacent cells, to identify distinct cell types or untangle complex cellular interactions. We employed this approach to investigate the zebrafish liver architecture and its similarity with that of mammals. Previous studies reported that canaliculi, representing the apical membrane of hepatocytes, exhibit an intracellular topology by forming tube-like membrane invaginations within single hepatocytes (Figs. 3B, 4A; (Lorent et al., 2004; Sakaguchi et al., 2008). In mammals, the apical tubules are lined by membranes of two adjacent cells, thus displaying intercellular shared topologies (Fig. 4A;(Treyer and Musch, 2013). However, canaliculi topologies in the zebrafish liver have not been assessed at single-cell resolution. To determine canaliculi topologies, we combined *fabp10a:kalTA4*; *fraeppli-caax*^*cph4*^ injected with *cre* mRNA, which can discriminate the morphologies of adjacent hepatocytes, with an antibody staining for the canalicular multidrug resistance protein 1 (Mdr1) marking mature canalicular membrane at 5 days post fertilisation (dpf). Combining commercial and custom-made image analysis tools (see Methods), we developed an image analysis pipeline to extract and quantify canalicular topologies (movie S3). This revealed in addition to the known intracellular canaliculi (Fig. 4A,B) a substantial proportion of intercellular canaliculi, shared between the membranes of two or more hepatocytes (Fig. 4A,C,D; movie S3), demonstrating plasticity of canalicular topology in zebrafish. Altered canalicular topologies are also observed upon chronic liver injury in mammals (Clerbaux et al., 2019; Kamimoto et al., 2020), further supporting that canaliculus topology is generally more plastic than previously known, as well as possible common mechanisms of canaliculi formation between zebrafish and mammals.

**Fig. 4.**
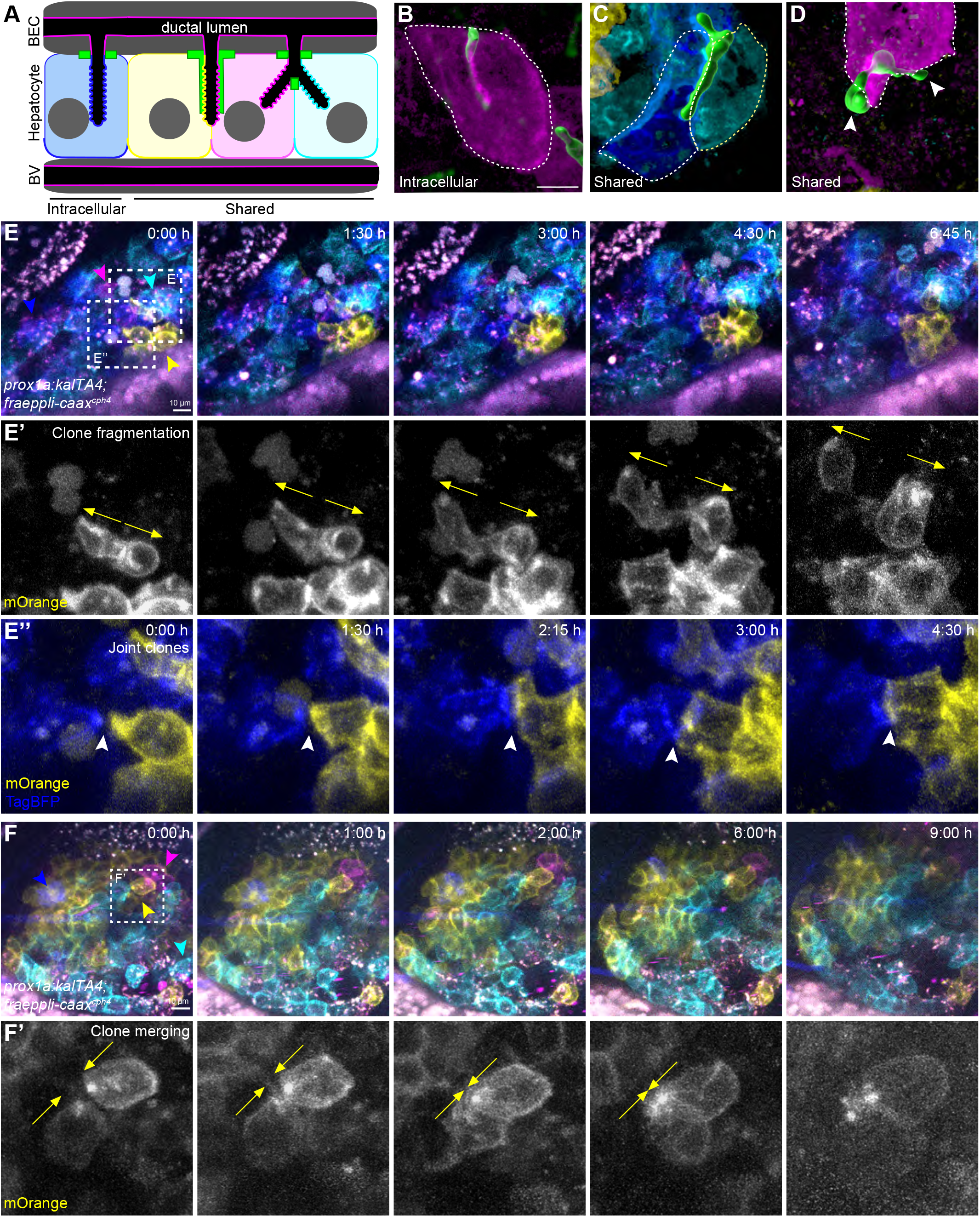
Topological analysis and long-term imaging of hepatocyte clones. (A) Schematic depicting intracellular and shared canalicular topologies. (B-D) Examples of canalicular topologies at 5 dpf scored by semi-automatic classification of FRaeppli-CAAX expressing hepatocytes, stained for canalicular protein Mdr1; arrowheads highlight canaliculi ends projecting into unlabelled hepatocytes; N=2; n=5 (see movie S3). (E-F’) *Live* imaging of 72 hpf *fraeppli-caax* embryos using sequential acquisition at 12-15 min intervals. Tracing dynamic cell rearrangement in the liver, reveals clone fragmentation as cells from the same clone move apart (E’; arrows), cells from two different clones aggregate into a joint clone (E”; arrowhead points to growing contact), or separated cells merge, forming new contacts (F’; arrows indicate direction of movement); N=3; n=10. (E,F) dashed frames indicate magnified areas shown in E’, E”, F’.

*Live* imaging is instrumental for elucidating morphodynamic cell behaviours driving organ and embryo development. Following the coordinated behaviour of several independent clones in the same embryo enables one to uncover the relative contributions of cell rearrangement and interactions during tissue formation. Sparse labelling with a single membrane-localised FP showed that directional progenitor migration determines liver position (Cayuso et al., 2016). However, it is unknown whether and to what extent cells rearrange as the organ differentiates. We monitored cell behaviours that occur as hepatocytes mature a functional morphology in *prox1a:kalTA4; fraeppli-caax* embryos injected with *phiC31* mRNA (movie S4). Larvae with clonal labelling in three or four colours in the liver were pre-selected and immobilized for about 12 hours of sequential image acquisition of 30-45 min intervals. Simultaneous tracking of individual clones over time showed continuous changes of clonal morphologies and interactions, including protrusive activity evident at clone edges or in small clones, indicating dynamic cell rearrangement and clone expansion during the phase of hepatic differentiation (Fig. 4E-F’). Tracing clone dynamics showed diverse behaviours, including cells of individual clones moving apart and intermingling with cells of adjacent clones, leading to clone fragmentation (Fig. 4E’; movieS4). Clone merging can occur in parallel, when cells that are initially separated, contact other clones, fuse and form tight clusters (Fig. 4E’’,F’). Combining real-time imaging with multicolour FRaeppli labelling thus revealed that differentiating hepatocytes are unexpectedly dynamic before they start to polarize, essential information for understanding the establishment of tissue architecture and organ growth (Rulands and Simons, 2016). It further highlights the suitability of FRaeppli for collecting time series of dense tissues at subcellular resolution, including those of internal organs. The presented applications of FRaeppli not only generate valuable insights into how complex tissues, such as the liver form, they also provide a glimpse at the extended experimental possibilities the tools offer.

In summary, the FRaeppli toolbox offers a new cell labelling system that is: (i) spectrally compatible with many existing transgenic reporter lines and cell labels, and (ii) amenable to sequential PhiC31 integrase-based cell labelling and conditional Cre/lox gene inactivation. Together with the growing number of conditional Cre-based gene inactivation tools (Carney and Mosimann, 2018) FRaeppli will enable connecting cell morphologies with gene function, which is the essential next step for understanding how tissues are built. The experimental versatility of FRaeppli thereby opens up powerful new applications for analyses of cellular behaviours in dense multicellular tissues.

## Acknowledgements

We apologise to colleagues, whose primary work was not included due to space constraints. We like to thank E. Caussinus and M. Affolter for sharing the Raeppli constructs, X. Guo for key cloning advice, R. Heyne for cloning of *fabp10a:kalTA4*, E. Ambrosio for help with processing of adult tissue. O. Andersson, W. Herzog, K. Koltowska, B. Hogan, C. Mosimann, D. Stainier, and D. Wilkinson for additional constructs and zebrafish lines. We thank M. Walther and K. Kristiansen in Zürich and the department of experimental medicine (AEM) in Copenhagen for expert fish care. We gratefully acknowledge the DanStem Imaging Platform (University of Copenhagen), the ScopeM (ETH-Zurich), and the Center for Microscopy and Image Analysis (University of Zurich) for their support & assistance in this work.

## Data availability

The authors declare that all data supporting the findings of this study are available within the article itself and its supplementary information files or from the corresponding authors upon reasonable request. The image data generated and analysed in this study are available from the corresponding authors upon reasonable request.

