## Supplemental Material for "FRaeppli, a multispectral imaging toolbox for cell tracing and dense tissue analysis in zebrafish"

### Materials and methods

#### Zebrafish husbandry

Zebrafish *(Danio rerio)* embryos and adults were kept according to standard laboratory conditions (Westerfield, 2000). Research was performed in accordance with ethical guidelines approved by the Danish Animal Experiments Inspectorate (Dyreforsøgstilsynet). The following transgenic lines were used: *tgBAC(prox1a:kalTA4)*$^{uq3bh}$ (Koltowska et al., 2015), *tg(hsp70l:gal4)*$^{fci1}$ (Cayuso et al., 2016), *tg(kdrl:EGFP)*$^{s843}$(Jin et al., 2005), *tg(hsp70l:cre)*$^{zdf13}$ (Feng et al., 2007), *tg(-2.8fabp10a:Cre,cryaa:Venus)*$^{s955}$ (Ni et al., 2012).

#### Constructs

*fraeppli-nls and fraeppli-caax*

NLS- and CAAX-tagged fluorescent proteins from *Drosophila* Raeppli were combined into a final pBlueScript (pBS) vector following the original strategy described in (Kanca et al., 2014). The order of FPs differs in the two FRaeppli cassettes: *pBS-FP-CAAX-cassette* contains mKate2-mTFP1-mOrange2-TagBFP whereas *pBS-FP-NLS-cassette* contains E2-Orange-mKate2-mTFP1-TagBFP. A fragment containing the SV40pA was amplified with primers #334 and #335 from the original pCS2+ vector (Rupp et al., 1994). The amplicon was cut with XbaI/SpeI and ligated into *pBS-FP-CAAX-cassette* and *pBS-FP-NLS-cassette*, both linearized with SpeI. A fragment containing the UAS promoter was amplified using primers #332 and #333 from *UAS:ephrinb1* (Cayuso et al., 2016) and cloned into pJET1.2 (Thermo Fisher). A cassette containing a minimal *attB* site, an *H2B-NeonGreen-STOP* coding sequence flanked by two *Lox2272* sites and several unique restriction sites for subsequent cloning steps (called ALNG), was synthesized as GeneBlock (IDT; sequence available upon request) and cloned using XbaI/NcoI into pJET-UAS cut with the same enzymes.

The H2B-NeonGreen contained in the *Lox2272*-flanked cassette, which was supposed to be expressed in unrecombined cells, was subsequently replaced with standard GFP amplified with primers #371 and #372 and cloned in the ALNG cassette by AscI/NdeI, given a suspected toxicity of NeonGreen in zebrafish. Unfortunately, neither version did express, probably due to a too far distance from the UAS.

The *phiC31 integrase* sequence followed by the *BGH polyadenylation (pA)* sequence was amplified with primers #351-340 from *pCDNA-phiC31-BGHpA* (Bischof et al., 2007) where a small sequence in the multiple cloning site was removed by XbaI/XhoI restriction and filled by Klenow DNA pol I (NEB). The full *phiC31 integrase* CDS was cloned after the ALGN cassette of *pJET-UAS-ALNG* using NsiI/BsrGI. The *cmlc2:mTurquoise* marker cassette was amplified from *UAS-self-cmlc2:mTurquoise* (derived from *pGEMT-cmlc2:mTurquoise*, kind gift from J. Goedhart) with primers #267 and #353 and inserted in pJET. This plasmid was linearized with NheI and the cassette containing *UAS-ALGN-phiC31 integrase* was cloned into it after release with AvrII/SpeI.

The resulting *pJET-UAS-ALGN-phiC31-cmlc2:mTurquoise* was used in subsequent steps to derive the final FRaeppli 1.0, which unfortunately could not be employed for the generation of transgenic lines, since the plasmid had already recombined in bacterial cells due to bacterial activation of the *UAS promoter*, followed by the unanticipated expression of the integrase. Details regarding various steps and different strategies attempted for solving the issue are available upon request.

To generate an intact FRaeppli 2.0 plasmid, refractory to recombination in bacterial cells, an intron from SV40 (Reddy et al., 1979) was placed at position 546 of the *phiC31 integrase* coding sequence, leading to a premature stop codon, which blocks the translation of a functional integrase only in bacterial cells. In eukaryotic cells the intron is spliced-out, therefore a functional integrase can be generated upon *UAS* activation. A cassette containing the second *lox2272* site, the first part of *phiC31 integrase*, the *SV40 intron* and linkers needed for subsequent cloning steps were synthesized as a GeneBlock (IDT; sequence available upon request) and subcloned into pJET-*fraeppli2.0*. Subsequently, the NdeI/BstEII fragment of pJET-*fraeppli2.0* cloned into *pJET-UAS-ALNG-phiC31-cmlc2:mTurquoise* also cut with the same enzymes, introduced the SV40 intron in the integrase CDS. The resulting plasmid was then cut with NotI and the fragment (*UAS-ALGN-phiC31-intron-cmlc2:mTurquoise*) cloned into *pBS-FP-CAAX-cassette* and *pBS-FP-NLS-cassette*. The *miniTol2* sequences (Urasaki et al., 2006), taken from *UAS-self- cmlc2:mTurquoise* were inserted in the resulting pBS vectors by substitution of the XhoI-bound fragment, generating the final FRaeppli2.0-NLS and FRaeppli2.0-CAAX, used for transgenesis.

All primers used in this study are listed in Table S1.

*fabp10a:kalTA4;cryaa:Venus*

The optimized Gal4 activator *kalTA4* coding sequence (Distel et al., 2009) was inserted after the *fabp10a* promoter (Her et al., 2003) into the *I-SceI-fabp10a:cre-cryaa:Venus* plasmid (Feng et al., 2007) where *cre* was removed with ClaI/NotI. An unnecessary *polyA* sequence was then removed by cutting with XbaI. The final plasmid was sequenced with primers #327 and #331.

*hsp70l:phiC31*

The EcoRI/NheI *phiC31-integrase-BGHpA* fragment was sub-cloned from the *UAS:phiC31-cryaa:Citrine* plasmid (this work) into the I-SceI containing backbone of the *fabp10a:kalTA4-cryaa:Citrine* plasmid, previously linearized with EcoRI/XbaI.

The *hsp70l* promoter was amplified from *hsp70l:mCherry-2A-wnt2* (Poulain and Ober, 2011) using primers #613 and #614, cut with ClaI/NsiI, and subsequently cloned into *I-SceI-phiC31 integrase* cut with the same enzymes. Primers #86, #382, #347 were used for sequencing the *I-SceI-hsp70l:phiC31 integrase* plasmid. The *he1a:palm-Citrine* marker cassette was generated by replacing the promoter sequence in the *UASself-cryaa:Citrine* plasmid (Cayuso et al., 2016) with the *hatching enzyme 1a (he1a)* promoter sequence (kind gift from the R. Koester laboratory). The *he1a:palm-Citrine-SV40polyA* cassette was amplified with primers #617 and #618 and cloned into I-SceI-phiC31 cut with Acc65I/SwaI, using blunt-end cloning.

*hsp70l:phiC31-sfGFP-NLS*

The previously *described I-SceI-hsp70l:phiC31-he1a:palm-Citrine* was modified by replacing the stop codon after the *phiC31* sequence with a linker sequence (5'-acttggttccag-3'). The resulting plasmid was called *I-SceI-hsp70l:phiC31-linker-he1a:palm-Citrine*. The *sfGFP* sequence (kind gift from H. Knaut laboratory) was amplified with primers #628 and #629 and cloned into *I-SceI-hsp70l:phiC31-linker-he1a:palm-Citrine* with PspOMI. Finally, the *NLS* sequence was added in frame at the C-terminal of the CDS by an intermediate subcloning step into *pBSII-BFP-NLS*. The final plasmid *I-SceI-hsp70l:phiC31-sfGFP-NLS-he1p:palm-Citrine* was sequenced with primers #382, #355, #581 and #615.

**Transgenesis**

*fraeppli2.0-nls* and *fraeppli2.0-caax* plasmids were injected into one-cell stage embryos together with *tol2* mRNA (30 pg mRNA and 20 pg DNA/embryo) as previously described (Kawakami et al., 2000). Individual F1 *fraeppli* fish were tested for recombination efficiency, using *cre* or *phiC31 integrase* mRNA injection. The three/four best lines are maintained as stable lines: *fraeppli-nls*$^{cph1\text{-}3,cph9}$ and *fraeppli-caax*$^{cph4\text{-}6}$. The *I-SceI-hsp70l:phiC31-he1a:palm-Citrine*, and *I-SceI-fabp10a:kalTA4-cryaa:Venus* plasmids were injected in one cell-stage embryos together with I-SceI enzyme (NEB) as previously described (Soroldoni et al., 2009). Stable transgenic insertions are maintained as *hsp70l:phiC31*$^{cph7}$ and *fabp10a:kalTA4*$^{cph8}$.

**FRaeppli activation**

To either prime or activate FRaeppli recombination, *cre* mRNA (20-25 pg) or *phiC31 integrase* mRNA (30-40 pg) were injected into one-cell stage embryos, as described (Felker and Mosimann, 2016; Mosimann et al., 2013). For temporally controlled activation, *fraeppli* and desired Gal4/KalTA4 driver lines were crossed to either *tg(hsp70l:cre)*$^{zdf13}$ or *tg(hsp70l:phiC31-he1a:palm-Citrine)*$^{cph7}$. Embryos were subjected to a 30-60 min heat-shock at 39°C and selected for recombination at desired stages. The recombination in Cre-primed embryos follows the sequential onset of transgenic Gal4/KalTA4 and subsequent FRaeppli-inbuilt PhiC31 integrase expression.

**HCR analysis**

mRNA expression of *hsp70l:phiC31* was analysed by in situ Hybridization Chain Reaction (HCR) RNA FISH version 3.0 (Choi et al., 2018). HCR probes against PhiC31 were purchased from Molecular Instruments, Inc. Fixation, detection, and amplification were performed according to the HCR v3.0 protocol for whole-mount zebrafish larvae (Choi et al., 2016), apart from the following modifications: 1 pmol of the probe set were used for detection, and only 15 pmol hairpin solutions were used for signal amplification. Samples were counterstained with 1:500 DAPI overnight at 4 °C. Embryos were mounted in Vectashield and imaged with a Zeiss 880 confocal microscope using a 40x oil objective. Maximum intensity projections of z-stacks were generated in Imaris and representative images are shown in the figures.

**Imaging**

Embryos and larvae intended for imaging experiments were raised in E3 medium (5 mM NaCl, 0.17 mM KCl, 0.33 mM $CaCl_2$, and 0.33 mM $MgSO_4$) supplemented with 0.2 mM 1-phenyl 2-thiourea (PTU, Sigma-Aldrich) to inhibit pigmentation. For *live* imaging, embryos and larvae were embedded in 0.8% low melting agarose and anesthetized with Tricaine (164 mg/L; MS-222, Sigma-Aldrich) dissolved in E3/PTU. For the time series in Fig. 4, four embryos in a single plate were concurrently imaged every 12-15 min for 8-12 hours using sequential acquisition parameters (Fig. S4A). Stained larvae were embedded in VectaShield (Vector labs) for whole-mount imaging. Whole liver lobes or vibratome sections of adult livers were cleared and imaged in SeeDB2S or embedded and imaged in VectaShield. Imaging was performed on Zeiss LSM 780, 880 and 980 confocal microscopes equipped with PMT detectors for sequential imaging and a spectral detector (GaAsP-PMT, 32 channels, 410-694 nm range, 8.9 nm bandwidth) for spectral acquisition. 5-colour sequential imaging was performed with a Leica Stellaris confocal microscope equipped with a tunable white-light laser and four PMT/HyD detectors. In spectral acquisitions, all four FRaeppli FPs were simultaneously excited with 405 nm, 456 nm and 561 nm lasers. In sequential acquisition experiments 515 nm and 594 nm excitation lasers were needed to discriminate between mOrange/E2-Orange and mKate2. Additional FPs or fluorophores were excited with a 488 nm and a 633 nm laser. Details of imaging parameters for different microscope set-ups are described in Fig S4A.

**Immunostaining**

Embryos were fixed with 4% PFA at 4°C overnight and stained as previously described (Thestrup et al., 2019). Adult fish were fixed overnight in 4% PFA at 4°C, using a rotator. Livers of adult fish were dissected out, embedded in 4% agarose/water, cut into 130 μm-thick sections using a vibratome (Leica) and stained with desired antibodies. Alternatively, dissected liver lobes were stained as whole-mount and subsequently cleared following the SeeDB2S protocol (Ke et al., 2016). The following primary and secondary antibodies were used: α-2F11 (mouse monoclonal, 1:10000, gift from Julian Lewis), α-MDR1 (rabbit polyclonal antibody, 1:2000, Santa Cruz, sc-8313), goat-anti-mouse 488 (1:500, Jackson Immunoresearch), goat-anti-rabbit 647 (1:500, Jackson Immunoresearch). Fixed embryos and adult liver lobes were incubated in PBS-TritonX 0.1%, Alexa Fluor 488 Phalloidin (1:1000, Molecular Probes, A12379) and TO-PRO-3 stain (1:50000, ThermoFisher) for 1 hour at room temperature or overnight at 4°C. Fresh samples were used for staining and imaging, since FRaeppli endogenous fluorescence is stable in fixed tissue for at least 1-2 weeks.

**Image analysis**

Images were processed with Bitplane Imaris and Zeiss ZEN software. Maximum intensity projections of z-plane stacks are shown, unless indicated differently in the figure legends. For FRaeppli CAAX images, the gamma value of each FRaeppli channel was increased to 1,2-1,5 to better visualize dark regions of the cell membranes. The gamma value was not altered for analysis purposes.

<u>Colour unmixing</u>

Lambda scans were spectrally unmixed using HyperSpectral Phasor software (HySP) (version 0.9.10) (Cutrale et al., 2017)(https://bioimaging.usc.edu/software.html). 16-bit spectral images stored in lsm5 format were opened in HySP with harmonic number 1. Images were denoised in the phasor window to reduce spectral scattering by 10-fold filtering. The minimum intensity values were manually adjusted to subtract dark background, and the phasor was displayed in logarithmic scale. Different clusters in the phasor plot corresponding to the spectra of the fluorescent proteins were manually gated using the polygon ROI selector (see Fig 3A') following the Detailed Guide to HySP (https://bioimaging.usc.edu/software.html#instructions). The overlap of the ROI in the image window was used to verify the ROI positions.

Of note, for *fraeppli-caax* we recommend comparing the sequential and spectral acquisition for any given experiment, since faithful colour assignment at cell interfaces of touching cells using the hyperspectral phasor plot is more complex. This also depends on the labelling density obtained. The most reliable, though slower option is to use sequential acquisition settings. This does not apply to *fraeppli-nls* acquisitions since nuclear signals never spatially intersect.

*Quantification of colour distribution*

To quantify colour choice in *fraeppli-caax*, embryos of the genotype *fraeppli-caax*$^{cph4}$*; prox1a:kalTA4*$^{uq3bh}$*; hsp70l:cre*$^{zdf13}$ were either injected with *cre* mRNA at one-cell stage or *hsp70l:cre* expression was induced at 33 hpf. Stacks of the first 30 $\mu$m of liver tissue were acquired by *live* imaging with spectral acquisition at 5 dpf, followed by manual annotation of the recombined cells in the Zeiss ZEN software. Embryos of genotype *fraeppli-nls*$^{cph1}$*; prox1a:KalTA4*$^{uq3bh}$; *hsp:phiC31*$^{cph7}$ were used to quantify colour choice in *fraeppli-nls*. Recombination was induced by heat shock at 36 hpf as described above. The left liver lobe of 5 dpf larvae was acquired *live* by spectral imaging. Sequential HySP unmixing was followed by segmentation in Imaris and determination of the number of nuclei using the Spots function.

*Quantification of FRaeppli transgene recombination*

DNA was extracted from single embryos of the indicated genotypes (Fig. S3F,G). PCRs amplifying recombined transgene species or un-recombined controls were performed (strategy in Fig. S3E). The following primers were used: #666-#667 (a; detecting a common region in the transgene), #170-#662 (b; detecting TagBFP in recombined transgene), #170-#346 (c; detecting mTFP1 in recombined transgene), #170-#663 (d; detecting mOrange2 in recombined transgene) and #170-#664 (e; detecting mKate2 in recombined transgene). The same PCR reactions were performed on 10 ng of FRaeppli 1.0 and 2.0 plasmid as control templates, since FRaeppli 1.0 recombines randomly during E. coli growth and thereby was used as positive control of recombination (see 'Constructs' section). Pictures of agarose gels were taken with different exposure times depending on the PCR, to avoid overexposing bands. Band quantification was performed with GelAnalyzer 19.1 (www.gelanalyzer.com; by Istvan Lazar Jr., and Istvan Lazar Sr.), using a morphometric algorithm to automatically define the background. Each band for the recombined versions (b-c) was first normalized to the intensity of the common band (a) from the same sample to correct for differences in DNA content. Then a second normalization factor for correcting the differences in picture exposure time was calculated by measuring the 1000 bp band of the ladder (GeneRuler 100bp plus, ThermoFisher) in each picture and applied to each band. Finally, the PCR efficiency, for detecting each recombined colour, was calculated by comparing the normalized intensities of the bands (b-e) of the positive control (FRaeppli 1.0 construct). The PCR-efficiency normalization factor for each colour was then applied to every experimental band of the corresponding colour.

*Analysis of hepatocyte canaliculus topology*

In 5 dpf stained larvae, FRaeppli FPs were imaged with spectral acquisition and the MDR-stained canaliculi (Alexa 488) in a separate acquisition with a PMT detector, since only a weak signal was detected in the spectral image. The spectral image was processed with the hyperspectral phasor software (see above) and seven channels were identified: four FRaeppli colours (B, C, Y, R) and the intersecting channels between Tag-BFP and mTFP1 (B/C), between mTFP1 and MDR (C/MDR) and between mOrange2 and MDR (Y/MDR). The intersecting channels were used in the analysis to correctly assign pixels that were positive for two adjacent colours in the spectrum, indicating spatial colocalization. The optimized unmixing gates were applied to all images and minimally adjusted to include all significant pixels in each image.

Segmentation was performed in Imaris. First, the seven unmixed channels (exported from the Phasor software) and the MDR channel of each image were combined in a single file in Imaris, following the application of a 5x5x5 median filter to the MDR channel. Subsequently, all the eight channels (cell membranes and canaliculi) were segmented in 3D with the 'Surface' function. An additional surface was created by merging the surfaces of the seven FRaeppli channels (cell membranes). The merged surface was used to select a subset of canaliculi overlapping with FRaeppli-labelled cells, by intersecting the two surfaces (FRaeppli membranes and canaliculi). The individual surfaces of the segmented membranes were subsequently used to create masked channels, whereby each colour-positive pixel was set to the same maximal value by thresholding (each included pixel=1; each excluded pixel=0). By doing this, intensity quantifications for each masked channel would also translate in relative volume estimation. For each FRaeppli-positive segmented canaliculus, several parameters were exported for further analysis: the Intensity Mean of each masked channel, indicating the relative volume of each canaliculus that was positive for that colour, the Shortest distance to Surface, indicating all the reciprocal distances between the canaliculi, and the Sphericity.

All parameters were imported into a custom-made script coded in R (Team, 2019; Team, 2020) to extract the topological information for all analyzed canaliculi. First, each canaliculus was associated with one or multiple colours, using a 5 % threshold of pixel occupancy for each channel, and the intersecting channels as quality controls. Second, the sphericity (<0.75) and distance (<0.5 um) between each canaliculus were used to classify canaliculi with an 'acinar' configuration, namely those shared between neighbouring hepatocytes which also contain an intracellular portion (Fig. 4A). Acinar canaliculi with >30% of pixels occupied by one or multiple colours were considered in the final classification.

A threshold of total colour occupancy of >70 % was applied for all the remaining canaliculi, which were subsequently divided in 'intracellular' or 'shared' by the presence of one or multiple colours, respectively. An additional distance filter was applied, to exclude intracellular canaliculi of neighbouring cells of the same colour in a 14 μm radius, the typical size of a hepatocyte, since shared or intracellular configurations cannot be discriminated in those cases. Thereby, only canaliculi included in isolated cells or cells surrounded by neighbours of different colours were included in the final classification. Additional information on the analysis and the script are available upon request.

### Supplemental Figures

**Fig. S1. Schematics of FRaeppli activation approaches.** (A) Overview of the FRaeppli construct. Starting from the left: *5xUAS* can control expression of the selected FP, as well as the FRaeppli-inbuilt PhiC31 integrase. The following *attB* site can recombine with one of the four *attP* sites upstream of each FP. A STOP cassette flanked by two *lox2272* sites prevents uncontrolled expression of the *phiC31 integrase* gene, downstream of the *UAS*. Next, *cmlc2:mTurquoise* serves as genetic marker for the FRaeppli transgene and separates further the *attB* from *attP* sites, thereby ensuring a minimal bias of transgene selection. The *phiC31 integrase* gene is expressed after Cre-mediated excision of the STOP cassette. Lastly, the four FP genes, each equipped with an upstream *attP* site and a 3' STOP codon, are arranged in a regular fashion and conclude with a single polyadenylation signal. (B) 2-step activation of FRaeppli colour selection: (i) Cre removal of the STOP cassette primes the FRaeppli cassette for recombination. Gal4 binding to the *5xUAS* activates the expression of *phiC31 integrase* gene, which (ii) can in turn recombine *attB* with one of the *attP* sites, self-excising the fragment in between, including the *phiC31 integrase* gene itself. (C) The addition of and exogenous source of PhiC31 integrase circumvents the need for Cre-medicated cassette excision and triggers *attB-attP* recombination in a one-step action. (D) Following colour selection, Gal4/UAS initiates the expression of the closest FP. Each FP gene ends with a stop codon, preventing expression of the remaining FPs, distal to the first one.

**Fig. S2. Characterization of FRaeppli-CAAX lines.** (A-C) representative images of three independent *fraeppli-caax* lines activated by *cre* mRNA injection (A) or heat-shock-driven *cre* expression (B,C). (D) Quantification of clonal abundance in the three independent transgenic *fraeppli-caax* lines shown in A-C. *fraeppli-caax*$^{cph4}$ exhibits the highest recombination efficiency, leading to the presence of >10 clones per field of view in 37.5% of embryos. Transgenic lines *fraeppli-caax*$^{cph6}$ and *fraeppli-caax*$^{cph5}$ give rise to sparser labelling. Number of quantified embryos per condition is indicated in each column. (E) Quantification of colour selection frequency in *fraeppli-caax*$^{cph4}$ livers (N=4; n=14).

**Fig. S3. PhiC31integrase recombines all four FRaeppli colours synchronously**. (A-C) HCR staining showing *phiC31 integrase* mRNA expression in *hsp70l:phiC31* embryos 2 and 6 hours after heat-shock induction (A,C), compared to controls without heat-shock (B). Maximum intensity projections of confocal z-stacks of the trunk region. As expected, the HCR signal is nuclear at 2 hours and more cytoplasmic after 6 hours, since the mRNA molecules have already been exported from the nucleus. (D) TagBFP fluorescence can be detected prior to the other FPs in *fraeppli-caax;hsp70l:gal4* embryos injected with *phiC31 integrase* mRNA. *Live* imaging shows tagBFP signal already at 22 hpf. (E) Schematic representations of the genomic *fraeppli-caax* construct before and after PhiC31 Integrase-mediated recombination. (F-G) PCR on genomic DNA from 10 and 24 hpf *fraeppli-caax*$^{cph4}$ single embryos activated by *phiC31 integrase* mRNA injection. (F') Primers 'a' amplify a common region of the FRaeppli construct, indicating the presence of the integrated transgene. Primer combinations 'b' and 'c' amplify only specific recombined forms of the FRaeppli transgene, when TagBFP (F'') or mTFP1 (F''') assume the first position after the promoter (schematic in E). Recombination for both colours is already detectable at 10 hpf detectable in all FRaeppli transgenic embryos. (G) PCR quantification of the recombined *fraeppli-caax* locus for all four colours, using the primer combinations indicated in (E). Recombination was triggered at 24 hpf by heat-shock-driven Gal4 expression, triggering inbuilt *phiC31 integrase* expression. 24 hours later (at 48 hpf), recombination for all colours was detected in 9/10 analysed embryos at variable degrees without bias, indicating synchronous and random colour-recombination.

**Fig. S4. Imaging configurations for spectral and sequential acquisition.** (A) Sequential and spectral FRaeppli acquisition settings for the microscope set-ups used in this study. (B) Schematic showing the inverse behaviour of imaging speed and post-processing time when applying spectral and sequential imaging modes. (C) Spectral acquisition with 2-photon excitation using two wavelengths is sufficient to excite all four FPs expressed in the liver at 4 dpf from *fraeppli-nls;prox1a:kalTA4* injected with *cre* mRNA. (D) Sequential and (E) spectral acquisition followed by unmixing using hyperspectral phasor analysis show comparable detection outcomes for all FPs in liver sections from adult *fraeppli-nls;prox1a:kalTA4* zebrafish. Embryonic *cre* mRNA injection triggered FRaeppli recombination.

**Fig. S5. Advanced labelling approaches for FRaeppli with spectrally-distinct markers for visualizing tissue context.** (A,B) Maximum intensity projections of sequentially acquired liver tissue from *fraeppli-nls;prox1a:kalTA4* adults cleared using the SeeDB2 protocol (A), and *fraeppli-caax;prox1a:kalTA4* 5 dpf larvae (B). Beyond FRaeppli, tissue was stained for Actin (Phalloidin-488, green) and the nucleus (TO-PRO, red) to visualize tissue architecture at the cellular and subcellular level. Embryonic *cre* mRNA injection triggered FRaeppli recombination.

**Movie S1. Maturation dynamics of phiC31-sfGFP-NLS.** Time-series of *hsp70l:phiC31-sfGFP-nls*, which can be detected from around 2 hours after heat-shock and the expression peaks at 6 hours. Without heat-shock transgene expression is undetectable.

**Movie S2. FRaeppli-NLS labelling of adult tissues.** Endogenous FRaeppli-NLS fluorescence is well preserved with SeeDB2 clearing and detectable beyond 120 μm, while the fluorescent signal disappears after only 30 μm of depth without clearing.

**Movie S3. Stepwise analysis of canalicular topologies in 5 dpf zebrafish livers.** FRaeppli-CAAX combined with α-Mdr1 staining (Alexa 488) undergoes sequential segmentation and topological analysis of hepatocyte canaliculi in larvae (see also Supplementary Methods), including examples of distinct canalicular topologies.

**Movie S4. Time-lapse of dynamic liver cell rearrangement in a 72 hpf *fraeppli-caax* embryo.** A Tag-BFP cell (arrow at the beginning of the movie) loses contact with its clonal neighbour and moves 2-3 cell diameter away. Simultaneous labelling of cells in different colours allows studying cell movements in the cellular context.

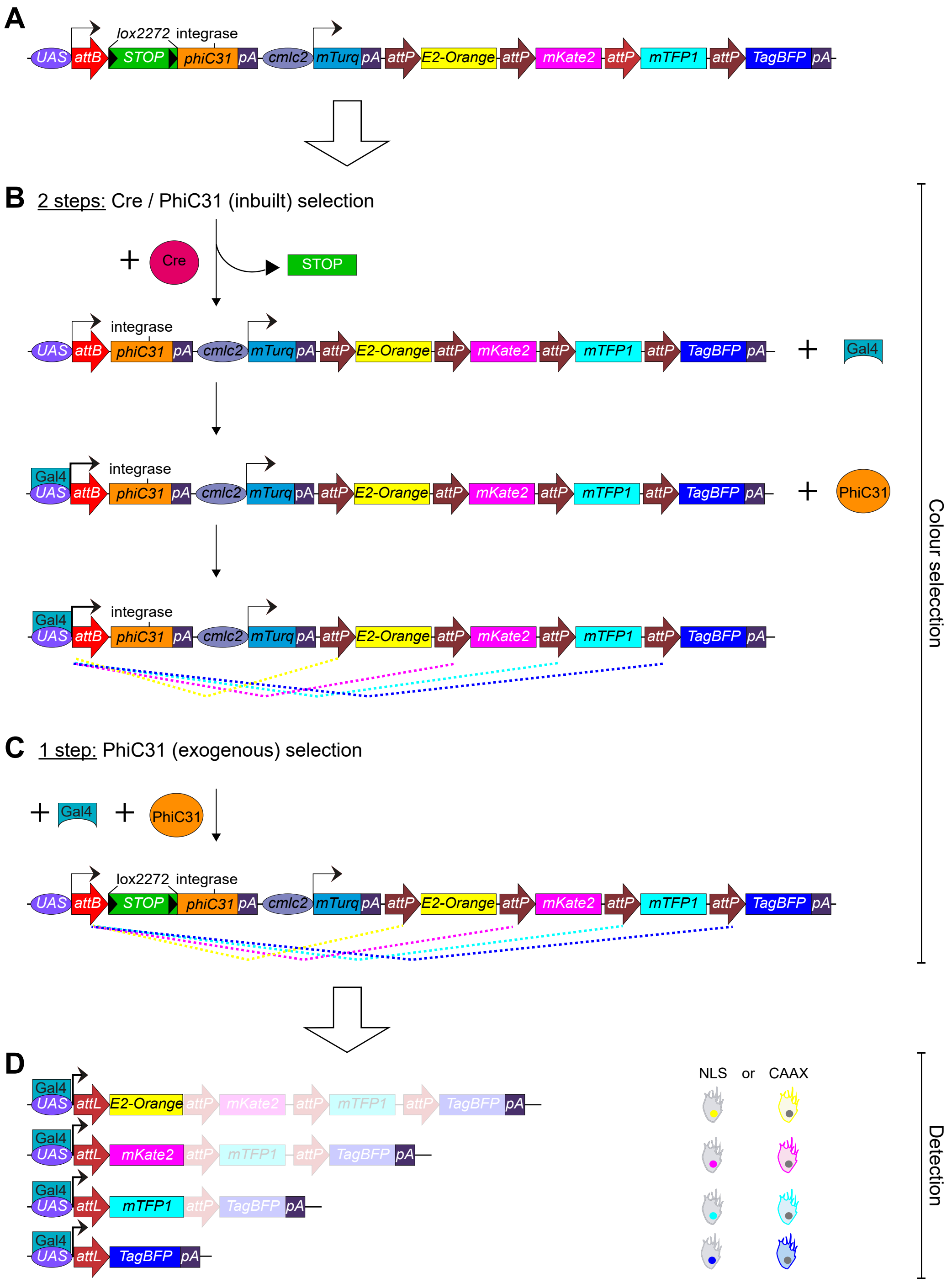


Figure S1 - Caviglia, Unterweger *et al.*

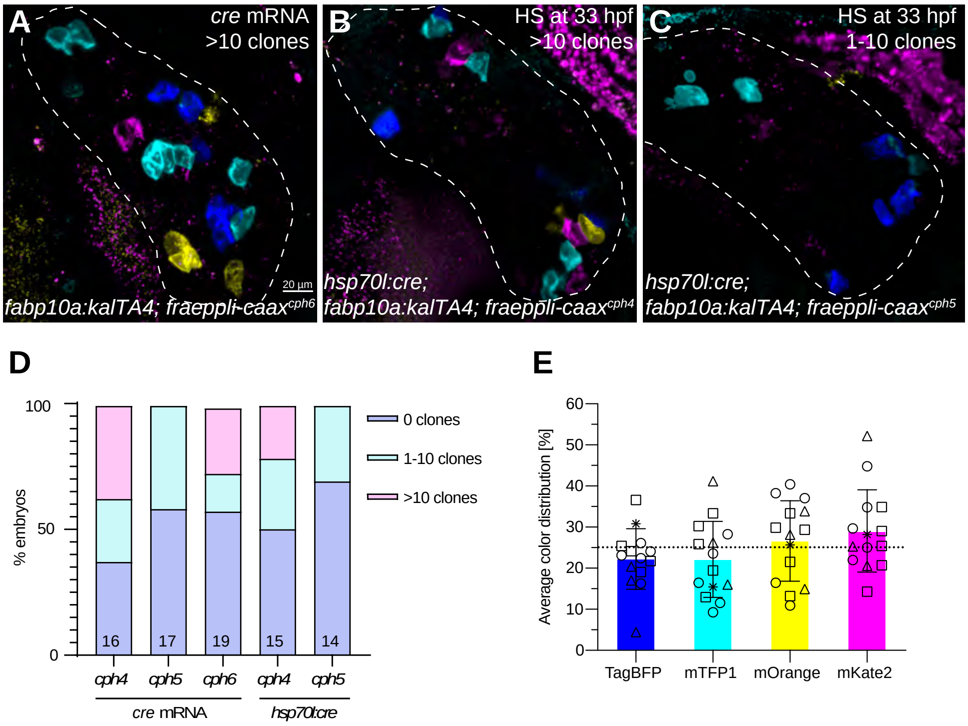


Figure S2 - Caviglia, Unterweger *et al.*

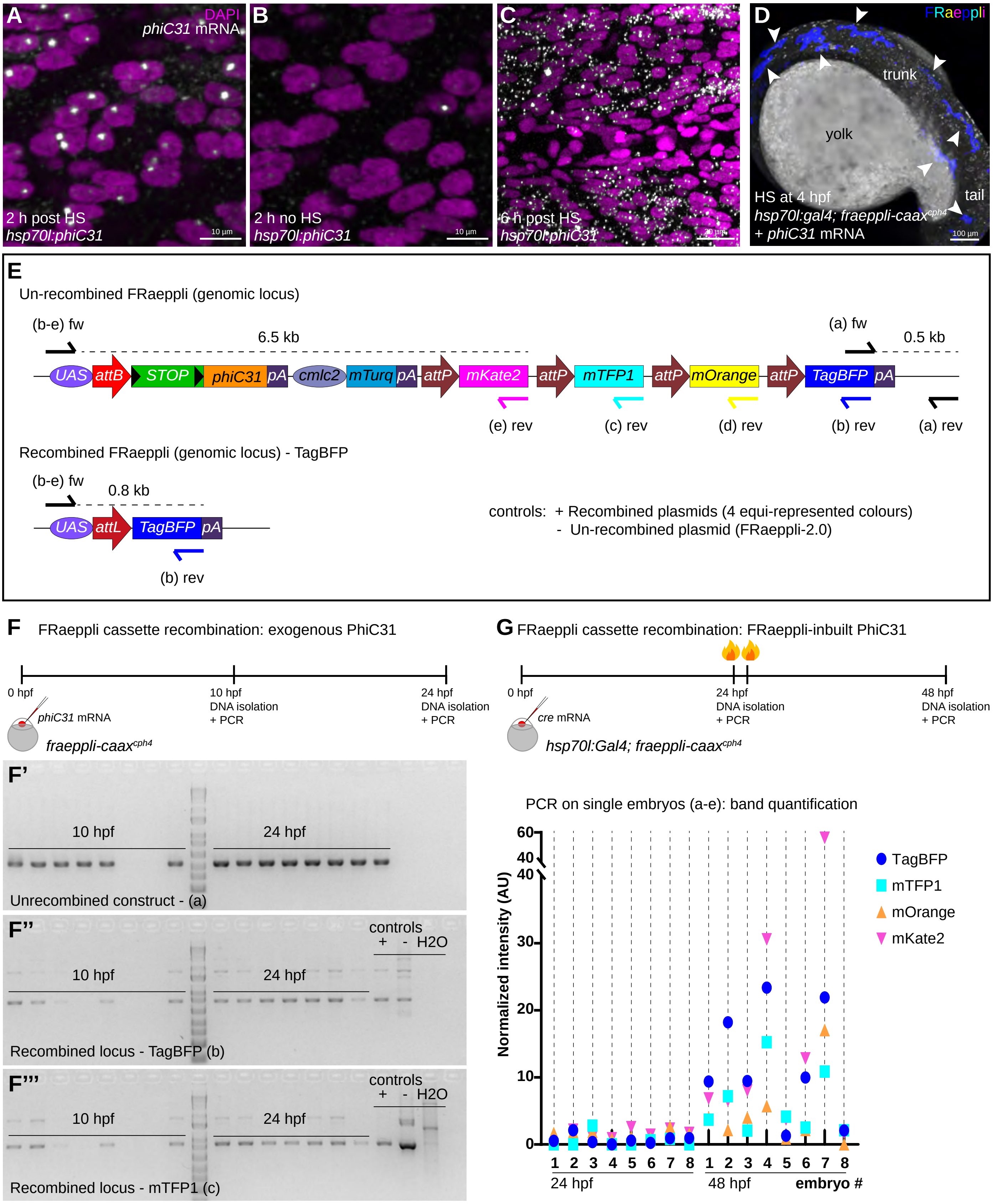


Figure S3 - Caviglia, Unterweger *et al.*

**A**

| | | | Sequential Acquisition | | | | Spectral Acquisition | |
|---|---|---|---|---|---|---|---|---|
| | | | Leica Stellaris 4 or 10 colors | Leica Stellaris 6 colors (live) | Zeiss LSM 880 6 colors (fixed) | Zeiss LSM 780 live timelapse | Zeiss LSM 780 | Zeiss Multiphoton |
| FRaeppli FPs | EXCITATION | TagBFP | 405 nm | 405 nm | 405 nm | 405 nm | 405 nm | 1100 nm |
| | | mTFP1 | 448 nm | 448 nm | 458 nm | 458 nm | 458 nm | 1100 nm |
| | | E2-Orange | 547 nm | 547 nm | 514 nm | 561 nm | 561 nm | 840 nm |
| | | mKate2 | 589 nm | 589 nm | 594 nm | 594 nm | 561 nm | 840 nm |
| | EMISSION | TagBFP | 415 - 460nm | 415 - 460 nm | 410 - 464 nm | 410 - 464 nm | Spectral Quasar detector* | Spectral Quasar detector* |
| | | mTFP1 | 477 - 553 nm | 477 - 487 nm | 463 - 579 nm | 464 - 499 nm | Spectral Quasar detector* | Spectral Quasar detector* |
| | | E2-Orange | 557 - 598 nm | 557 - 598 nm | 535 - 633 nm | 570 - 597 nm | Spectral Quasar detector* | Spectral Quasar detector* |
| | | mKate2 | 600 - 805 nm | 600 - 622 nm | 599 - 696 nm | 597 - 695 nm | Spectral Quasar detector* | Spectral Quasar detector* |
| additional spectra | EX | green / GFP | - | 488 nm | 488 nm | - | - | - |
| | | far-red | - | 642 nm | 633 nm | - | - | - |
| | EM | green / GFP | - | 513 - 546 nm | 520 - 575 nm | - | - | - |
| | | far-red | - | 660 - 790 nm | 643 - 747 nm | - | - | - |

* = 32 channels, 8.9 nm bandwidth, spectral range: 410nm - 694nm

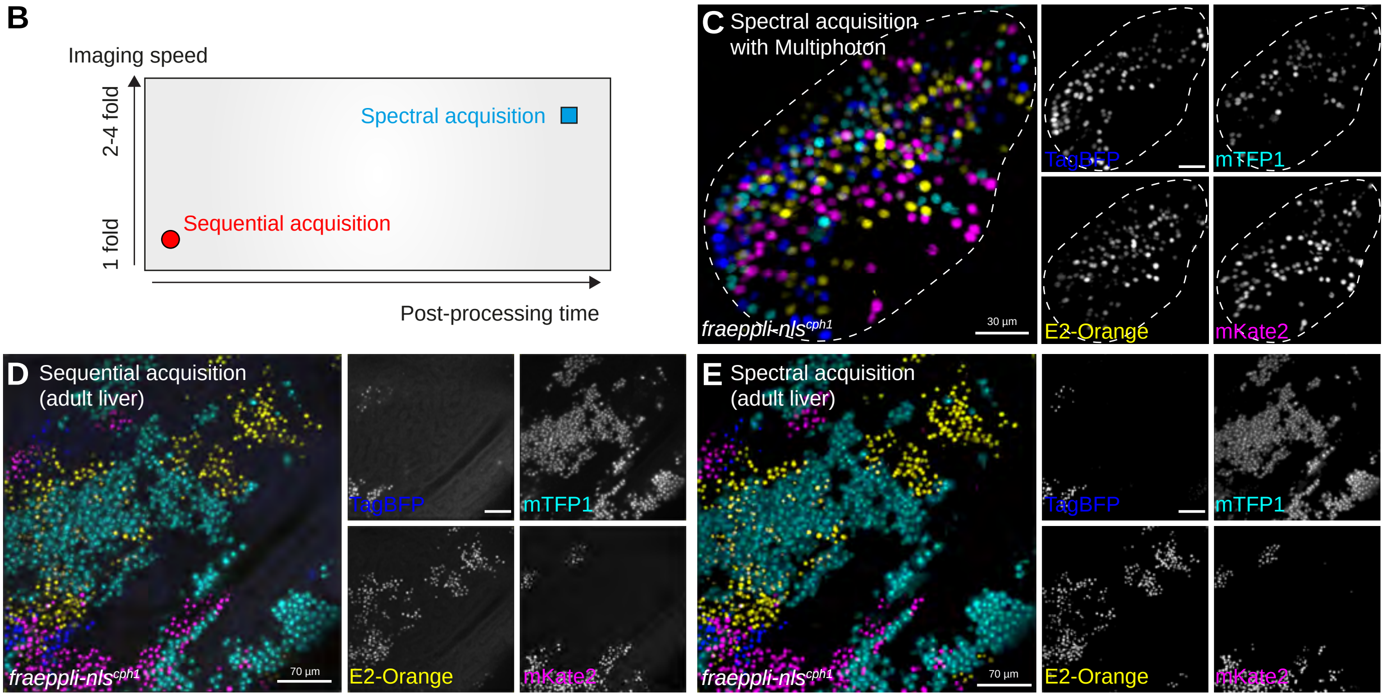


Figure S4 - Caviglia, Unterweger *et al.*

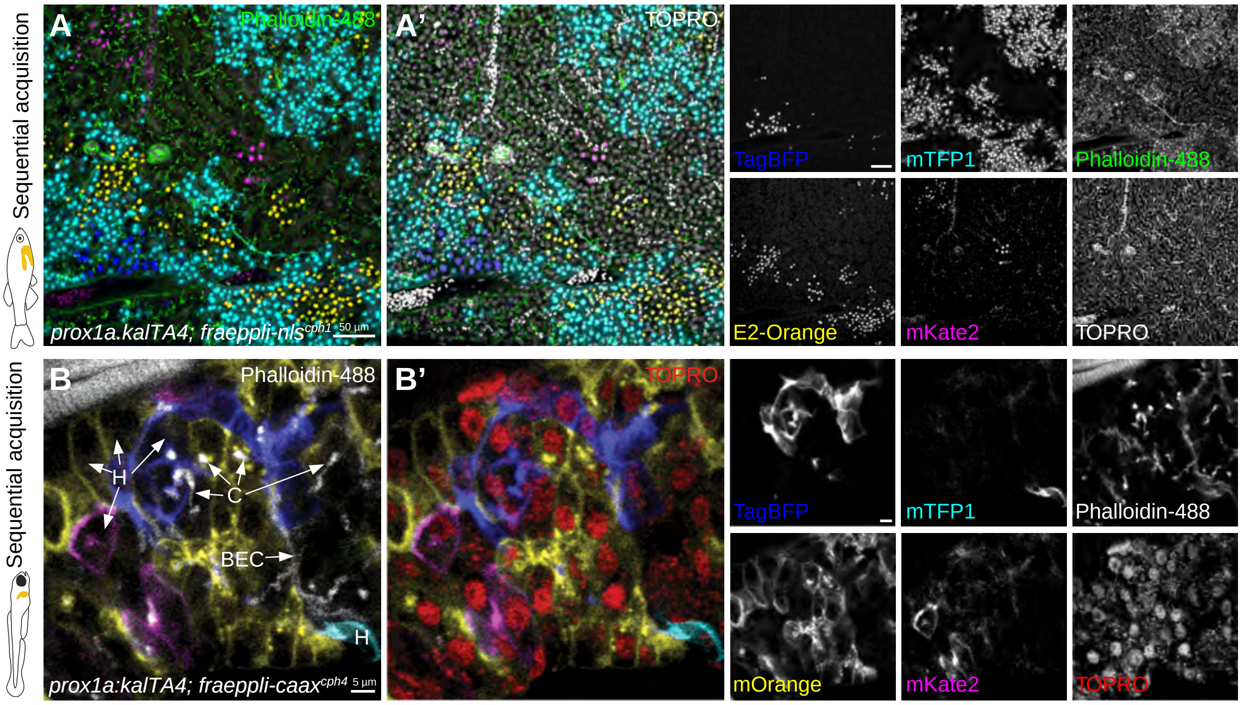


Figure S5 - Caviglia, Unterweger *et al.*

**Table S1**

| Primer # | Sequence |
|---|---|
| 86 | TAATACGACTCACTATAGG |
| 170 | CTGCAGGTCGGAGTACTGTC |
| 267 | AATTGAATTGCGCTGATGC |
| 327 | CACTTTGTGTTGAGCGG |
| 331 | TTGGTCGGTCATTTCGG |
| 332 | CTATCGGCCGCCTAGGTAGGGATAACAGGGTAATACCGGTC CTGCAGGTCGGAGTACTGT |
| 333 | ATCGAATTCGTGTGGAGGAG |
| 334 | CTAGTCTAGAAGTGAGTCGTATTACGTAGATCC |
| 335 | CTAGACTAGTTAGGGATAACAGGGTAATCGAATTAAAAAACCT CCCAC |
| 336 | TTAAGGAAGTAAAAGTAAAAGCAAGAA |
| 340 | TGTACAGGTTCTTTCCGCCTCAGAA |
| 341 | ACCTCCAGGTTGATGGTGTT |
| 342 | CCTGAAGGGCAAGATCAAGA |
| 343 | ACGGCACCTTCATCTACCAC |
| 346 | GTTCAGGATCTCGATGCGGTG |
| 347 | AAGAACATCGATTTTCCATGGCAG |
| 348 | TTTATACGAAGTTATCCTAGCATGCA |
| 349 | GTATCCTATACGAAGTTATTTACATATGC |
| 350 | GCGTGCCTGATCTTGTTGAA |
| 351 | CTATATGCATCAAGCAAACATGGACACGTACGCGGGTG |
| 353 | CTAGGCTAGCGCAGGCCTAAATCAGTTGTG |
| 355 | CCGGGATCACTCTCGGCAT |
| 356 | GACCAACAGCAAAGCAGACA |
| 371 | ATATGGCGCGCCTATGGTGAGCAAGGGCGAG |
| 372 | CTAGCTACATATGCTTGTACAGCTCGTCCATGCC |
| 382 | GAAATGCCCGACGAACCTG |
| 581 | CACATGAAGCAGCACGACTTCT |
| 613 | CTAATATCGATTTCTGCAGCCCGGGGGATC |
| 614 | ATACATGCATGCTCTAGAGGCTCGACTGC |
| 615 | CGCTGAGTGGTATGAGCTTC |
| 617 | CTAGGGTACCTCAACCACTCCAGGCATAGC |
| 618 | TATACCGCGGGATCTGGCCATCTAGAGCGG |
| 623 | TTATGGATCCCAAGCAAACATGGACACGTACG |
| 624 | ATATGGGCCCACTTGGTTCCAGCGCCGCTACGTCTTCCGTG |
| 628 | TTATGGGCCCATGGTGAGCAAGGGCGAGG |
| 629 | ATATGGGCCCTCAACGGGGAACCCGAAGC |
| 650 | ATATGGATCCGGGCCCATGGTGAGCAAGGGCGAG |
| 651 | ATATCTGCAGACGGGGAACCCGAAGCTC |
| 662 | TATATGTGGTCTTGATGTTTGC |
| 663 | ACTGCTCCACCACGGTG |
| 664 | GCTTCAGCCTCATCTTGATCTT |
| 666 | GGAGCCATCTGATCGCAA |
| 667 | TAGGGATAACAGGGTAATCGAATT |
